# Neonatal 6-OHDA lesion of the SNc induces striatal compensatory sprouting from surviving SNc dopaminergic neurons without VTA contribution

**DOI:** 10.1101/2021.03.26.437271

**Authors:** William Tanguay, Charles Ducrot, Nicolas Giguère, Marie-Josée Bourque, Louis-Eric Trudeau

**Affiliations:** Department of Pharmacology and Physiology and Department of Neurosciences, Faculty of Medicine, Central Nervous System Research Group (GRSNC), University of Montreal, Quebec, H4T 1J4 Canada

**Keywords:** Parkinson’s Disease, Dopamine, Vulnerability, Axonal Arborisation, 6-Hydroxydopamine, Substantia Nigra

## Abstract

Dopamine (DA) neurons of the *substantia nigra pars compacta* (SNc) are uniquely vulnerable to neurodegeneration in Parkinson’s disease (PD). We hypothesize that their large axonal arbor is a key factor underlying their vulnerability, due to increased bioenergetic, proteostatic and oxidative stress. In keeping with this model, other DAergic populations with smaller axonal arbors are mostly spared during the course of PD and are more resistant to experimental lesions in animal models. Aiming to improve mouse PD models, we examined if neonatal partial SNc lesions could lead to adult mice with fewer SNc DA neurons that are endowed with larger axonal arbors because of compensatory mechanisms. We injected 6-hydroxydopamine (6-OHDA) unilaterally in the SNc at an early postnatal stage at a dose selected to induce loss of approximately 50% of SNc DA neurons. We find that at 10- and 90-days after the lesion, the axons of SNc DA neurons show massive compensatory sprouting, as revealed by the proportionally smaller decrease in tyrosine hydroxylase (TH) in the striatum compared to the loss of SNc DA neuron cell bodies. The extent and origin of this axonal sprouting was further investigated by AAV-mediated expression of eYFP in SNc or ventral tegmental area (VTA) DA neurons of adult mice. Our results reveal that SNc DA neurons have the capacity to substantially increase their axonal arbor size and suggest that mice designed to have reduced numbers of SNc DA neurons could potentially be used to develop better mouse models of PD, with elevated neuronal vulnerability.

**Graphical abstract text:**

- We describe a technique to induce the loss of approximately 50% of SNc DA neurons in neonate mice using unilateral intranigral 6-OHDA (left panel).
- Compensatory axonal sprouting was observed in the striatum as early as 10 days following the lesion (at P15), with effects lasting until adulthood (P90).
- Conditional AAV-mediated expression of eYFP (green) reveals SNc DA neurons, projecting to the dorsal striatum (middle panel), and not VTA DA neurons, projecting to the ventral striatum (right panel), as the main source of compensatory axonal sprouting.

**Graphical abstract:** 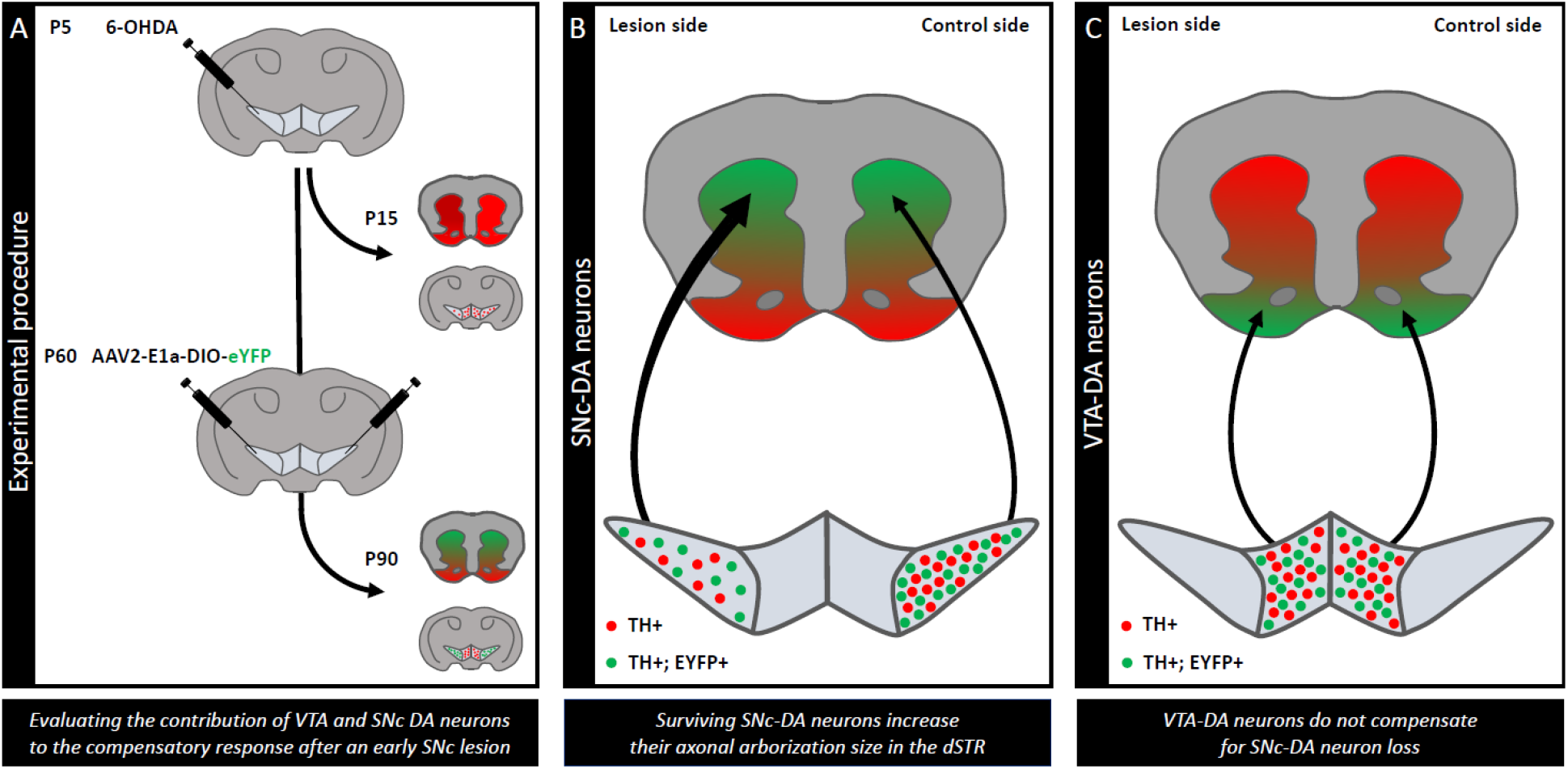

## Introduction

The lack of animal models of Parkinson’s disease (PD) faithfully reproducing the progressive and selective nature of neuronal loss observed in this disease in humans represents a major hurdle for current research efforts. While neurotoxin-based models, such as those using MPTP or rotenone, are useful to model some aspects of PD pathophysiology because they cause degeneration of substantia nigra pars compacta (SNc) dopamine (DA) neurons (Hamadjida et al., 2019; Segura-Aguilar, 2019), these models do not allow to reproduce well the progressive nature of cell loss and behavioral dysfunctions of this disease. In mice, toxins such as MPTP also lead to reversible downregulation of tyrosine hydroxylase (TH), complicating the quantification of definitive cellular loss (Mitsumoto et al., 1998). Repeated toxin administration regimens can be used as an attempt to better model the disease (Bové and Perier, 2012; Prediger et al., 2011), but these still fail to trigger self-sustaining neurodegeneration once the acute effect of the toxin dissipates.

Genetic models of familial forms of PD, deriving from the production of Parkin, Pink1, DJ-1 or LRRK2 knockout or knockin mice have provided disappointing results, with little if any spontaneous loss of SNc DA neurons and little if any observable functional or behavioral deficits (Dave et al., 2014; Kitada et al., 2007, 2009a, 2009b; Sanchez et al., 2014). However, such genetically manipulated mice have been reported to be more vulnerable to neurotoxins (Haque et al., 2012), and recently to infections (Matheoud et al., 2019).

Viral overexpression-based or fibril-based models of α-synuclein pathological aggregation and toxicity have shown some promise by successfully reproducing the progressive transmission of pathology (Ulusoy et al., 2010; Van Den Berge et al., 2019; Volpicelli-Daley et al., 2011; Yamada et al., 2004). However, concerns have been raised with regards to the supra-physiological nature of regionally concentrated administration of significant quantities of α-synuclein.

Irrespective of the model used, one recurrent concern with mouse models of PD has been the seemingly higher resilience of DA neurons in this species compared to humans. The origin of the intrinsically high vulnerability of nigral DA neurons is not completely understood, but the very large axonal arbor size of these neurons has been proposed to be a major determinant because of its tight link with mitochondrial function and associated oxidative stress (Franco-Iborra et al., 2015; Pacelli et al., 2015; Parent and Parent, 2006; Prensa and Parent, 2001). Because it is estimated that DA neurons in the mouse brain are endowed with a much smaller axon length and complexity compared to humans (Bolam and Pissadaki, 2012; Pissadaki and Bolam, 2013), this might make mouse DA neurons intrinsically more resilient compared to those of humans. In line with this observation, recent work showed that deleting the D2 autoreceptor gene from DA neurons leads to disinhibited axonal arbor growth and increased vulnerability to 6-OHDA (Giguère et al., 2019). Because lack of D2 receptor activation could influence vulnerability through other mechanisms in addition to axonal arbor expansion, complementary approaches to increase axonal arbor size in DA neurons are necessary. One such approach could be to exploit compensatory axonal sprouting mechanisms. In the adult brain, nigral DA neurons have been shown to possess a limited capacity for such axonal sprouting following various types of lesions (Robinson et al., 1994; Blanchard et al., 1996; Song et al., 2000; Deumens et al., 2002). However, neonate neurons in multiple regions of the nervous system have been previously shown to posses an extensive compensatory axon growth capacity during the first week following partial lesions (Filbin, 2006; Goldberg et al., 2002). Neonatal 6-OHDA lesions have been previously used to induce near complete loss of SNc DA neurons in the neonatal rodent brain and examine how striatal axonal sprouting by 5-HT neurons contributes to motor adaptations observed in these animals at the adult stage (Fernandes Xavier et al., 1994, Luthman et al., 1990, Luthman et al., 1987, Sivam et al., 1987, Snyder et al., 1986, Stachowiak et al., 1984, Stellar et al., 1988, and Towle et al., 1989). However, whether more partial lesions of SNc DA neurons could promote compensatory growth by surviving nigral neurons has not been examined previously. In the present work, we show that early postnatal chemical ablation of a subset of SNc DA neurons leads to substantial compensatory expansion of the axonal arbor of the surviving neurons in the striatum. This strategy shows great promise to further assess the hypothesis of axonal arbor size as a neuronal vulnerability factor in PD.

## MATERIALS AND METHODS

### Experimental subjects

All experiments were performed using male DATCre^+/−^mice on a C57/Bl6 background (Zhuang et al., 2005), in accordance with protocols approved by the Animal Experimentation Ethics Committee (CDEA) of the Université de Montréal. Mice were injected unilaterally at age P5 with a single intranigral dose of 6-OHDA and were sacrificed either at P15±1 or at P90±2. Some mice also received at P60 an intracerebral injection of AAV2-eYFP in the SNc bilaterally, or a single midline injection in the VTA.

### Unilateral injection of 6-OHDA in the SNc of P5 mice

The duration of all intracerebral injections was minimized, with an average surgical duration of 15 min. A solution of 6-hydroxydopamine (6-OHDA) was prepared less than 30 min before starting surgical procedures, at a concentration of 0.25mg/ml in normal sterile saline containing 0.02% ascorbic acid. A solution of 2.8mg/ml desipramine in normal sterile saline was prepared, aliquoted and kept at −80°C until use. Mice were given 30μl of desipramine subcutaneously 30 min prior to surgery. Mice were cryoanesthetized by being placed in a vinyl glove and submerged in ice and water for 100 to 120s and were then placed on a head mold allowing immobilization of the head, similar to the one described previously for intracerebral injections in neonate rats (Davidson et al., 2010; Sierra et al., 2009). This head support was then fixed on a stereotactic frame (Digital Lab Standard Stereotaxic for Mice with LED Display, Harvard Apparatus). The mold allowed the animal to be maintained at 4 to 5°C during the duration of surgery using a refrigerated block. A solution of 10% povidone-iodine was applied to the scalp and an incision was performed at the level of the lambdoid sinus. A 31-caliber beveled needle was fixed to the stereotactic system and lowered to the confluence of the lambdoid suture, which was used as a reference point. The needle was then moved to the (*x,y*) coordinates of the injection site and slowly lowered at these coordinates to pierce the skull and dura, after which the needle was slowly raised and replaced with the injection needle (caliber 32, non-beveled) joined by polypropylene tubing to a Hamilton injection syringe mounted on a microinjector. The injection needle was repositioned to lambda and coordinates were reset at that point, before displacing the needle to the (*x,y*) coordinates of the SNc for 6-OHDA injection (Lambda: 0,0,0, *x* (rostrocaudal) = −1,4 mm, *y* (mediolateral) =0,0 mm, *z* (dorsoventral) = −4,0 mm). The needle was then lowered to the *z* coordinate of the injection site over a period of 60s, with special care to avoid pulling on the meninges. Once the site was reached with the injection needle, 0.5μL of 6-OHDA was injected at a rate of 5μl/h, and the needle was then kept in place for 180s to allow proper diffusion of the bolus. The needle was then slowly withdrawn over a period of 60s and the wound was closed using surgical glue (Surgi-lock, 2-octyl-cyanoacrylate). A tissue sample was taken from the ear for genotyping and the animal was subsequently returned to the heated cage for termination of cryoanesthesia. Once all the animals were injected and had regained normal motor activity levels, they were returned to the parental cage until weaning (P21). Experimental cages containing injected animals were monitored at 24h and 48h following surgery without displacing the cage to minimize parental stress.

### Injection of eYFP virus in the SNc and VTA of P60 mice

DATcre^+/−^mice received an injection of rAAV2/Ef1a-DIO-eYFP virus (UNC GTC Vector Core, lot AV4842d, 4 × 10^12^virions/ml) at P60±2, either in the SNc bilaterally or in the VTA at the midline. Surgeries lasted from 30 to 60 min per animal, depending on the number of injection sites. The virus solution was diluted 1:4 in a 0.9% NaCl aqueous solution, and stored at −80°C. At the time of surgery, the virus aliquot was kept on ice until use during surgery. The stereotaxic system used was the same as for neonate injections of 6-OHDA (Harvard Apparatus), except that adult ear bars were used instead of the neonate head mold. The stereotactic frame was set up to accommodate a surgical drill and a Hamilton syringe allowing for manually controlled injections. A Hamilton syringe (Hmlt700, Sigma-Aldrich) (volume of 10μl, internal diameter of 0.485μm) attached to a glass micropipette, was filled with mineral oil and mounted to a stereotactic injector arm. The animal to be injected was deeply anesthetized using oxygenated air containing 3% isoflurane and was then placed on the stereotaxic frame with the head immobilized using the ear bars. Isoflurane (1.5-2%) was used during surgery for maintenance of anaesthesia. Eye drops were applied to the eyes of the animal to minimize risk of damage to the cornea. Hair was shaved on top of the animal’s head and the surgical site was cleaned with a 10% povidone-iodine solution. A solution of 1mg/ml of Marcaïne (bupivacaïne chlorhydrate) was injected subcutaneously under the scalp at a dose of 2μl/g. After a 60s pause, a 1cm skin incision was made sagittally using a scalpel, ending 3mm caudally to the interocular line, to expose the skull. Conjunctive tissue was scraped with the scalpel and the skull surface was washed with saline and dried. Under binoculars, the tip of the surgical drill (1mm bit) was placed at bregma and stereotaxic coordinates were set to zero. For the bilateral SNc injections, coordinates were: × (rostrocaudal) −3.3 mm, y (mediolateral) ±1.5mm, z (dorsoventral) −4.0mm, and injection volume was 0.8μl. For the central VTA injections, coordinates were: × (rostrocaudal) −3.1 mm, y (mediolateral) 0.0mm, z (dorsoventral) −4.5mm, and the injection volume was 0.4μl. A surgical drill was used to burr one or two holes, as required, taking care not to tear the underlying meninges. The drill was then withdrawn and replaced by the stereotaxic injector arm. The dura was then delicately pierced using a dural hook. After eliminating any air bubble in the glass micropipette, a volume of virus larger than the volume to be injected was pipetted on a piece of parafilm and aspirated in the injection micropipette. Under binoculars, the glass micropipette was then positioned at bregma, stereotaxic coordinates were set to zero and the tip of the micropipette was moved to the x,y coordinates of the desired injection site. The syringe was then slowly lowered to the z coordinates of the targeted structure over a period of 60s and a resting time of 120s was observed before starting the injection, which was made at 0.25μl/min. A resting time was observed for 5 min before withdrawing the micropipette tip over a period of 60s. If another injection had to be delivered to the second site, the injector wheel was turned a few times to check for micropipette clogging or damage, before reaching the x,y coordinates of the second injection site and making the second injection. At the end of the procedure, the scalp was hydrated with saline solution and the wound closed using 3 to 4 simple stitches (Ethicon®, Cornelia, GA, USA). A subcutaneous injection of carprofen solution (5mg/kg at 50mg/ml) was then administered postoperatively in a 0.9% saline solution bolus (0.1ml per 10g of body weight) to rehydrate the animal and provide postoperative analgesia. The animals were monitored until their return to wakeful state and were then put back in their cage with food pellets and a weighing boat containing Ensure® and were monitored at 24h and 48h after surgery.

### Animal perfusion and brain tissue preparation

Mice were sacrificed either at postnatal day 15±1, or 90±2. Animals perfused at P15 were deeply anesthetized with 0.25ml of chloral hydrate (24mg/ml) injected intraperitoneally and mice perfused at P90 were anesthetized using an intraperitoneal injection of pentobarbital (100mg/kg at a concentration of 7mg/ml). Animals were transcardially perfused with 100ml of phosphate buffered saline (PBS), followed by 100ml of 4% paraformaldehyde (PFA) solution. The brain was then extracted from the skull and preserved in a 4% PFA solution at 4°C. Brains were then transferred in a sucrose solution for a period of 48h at 4°C before being frozen. Subsequently, the brains were cut coronally using a cryostat in 40μm thick sections. Sections caudal to the anterior-most part of the striatum were serially distributed on a six-well plate in antifreeze solution. When mesencephalic sections containing the anterior-most SNc were cut, they were placed in a new series of six wells on a new plate.

### Immunofluorescence

For TH immunohistochemistry, a well was chosen at random amongst the six wells containing striatal sections for each animal of the experimental groups, to obtain a sampling of 1 in 6 sections for the whole striatum. Sections were then rinsed in 0.01M phosphate buffered saline (PBS) three times and incubated for 60 min on an agitation plate in a blocking solution (Triton X-100 0.3%, NaN3 0.02%, BSA 10.5%, goat serum 5%). Sections were then incubated during 24h in the primary antibody solution (anti-TH rabbit, Millipore, AB152(CH), 1:2000) (Triton X-100 0.3%, NaN3 0.02%, BSA 0.5%, goat serum 5%). After a triple-rinse in PBS, sections were incubated for 120 min in the secondary antibody solution (Alexa 488 goat anti-rabbit, A-11008, ThermoFisher Laboratories, 1:500), rinsed again three times in PBS, and mounted on microscope slides using 120μl of mounting medium (FluoromountG). For TH and eYFP double immunohistochemistry in P90 mice, the rabbit anti-TH antibody (1:2000) was detected using a goat anti-rabbit secondary antibody coupled to Alexa 546 (ThermoFisher Scientific, A-11010, 1:500) and eYFP was detected using detected using a chicken anti-GFP antibody (Aves Labs, GFP-1020, 1:2000) detected with an anti-chicken secondary antibody coupled to Alexa 488 (ThermoFisher Scientific, A-11039, 1:500).

### DAB immunohistochemistry and stereological counting

For stereological counting of mesencephalic TH positive neurons, one well amongst the six wells containing the mesencephalic sections was selected at random for each animal. Sections from this well were rinsed in 0.01M PBS for 10 min and then rinsed in 0.01M PBS containing 0.9% H2O2 for 10 min to inactivate endogenous peroxidase and rinsed again three times in 0.01M PBS (3×10 min rinsing time). Sections were then incubated at 4°C for 48h in the primary antibody solution (anti-TH rabbit, Millipore, AB152(CH), 1:1000) diluted in 0.01M PBS containing 0.3% Triton X-100 and 0.02% NaN_3_, and then rinsed three times in PBS. The sections were then incubated in the biotinylated secondary antibody solution (Biotin-SP goat anti-rabbit, Jackson ImmunoResearch Laboratories, 111-065-045, at 1:200 in 0.01M PBS containing 0.03% Triton X-100). After three 10-min rinses in PBS, sections were incubated for 3h at room temperature in the streptavidin horseradish peroxidase solution (Sigma, GERPN1231, diluted to 1:200 in 0.01M PBS containing 0.03% Triton X-100). Following three 10-min rinses in PBS and a 1 min rinse in acetate buffer 0.1M, sections were transferred for 1 min in wells containing the DAB reaction solution (DAB 1mg/ml, glucose oxidase 8mg/ml, Nickel ammonium sulfate 50mg/mL, glucose 2mg/ml, in acetate buffer 0.1M). Sections were immediately rinsed for 10 min in an acetate buffer 0.1M, mounted on microscope slides and left to dry for 72h. To defat sections and proceed with counterstaining, slides were submerged in successive baths of water (2×2min), cresyl violet solution preheated to 37°C (1×2min), ethanol (50% 1×1min, 70% 1×1min, 90% 1×1min, 100% 2×1min), isopropanol (1×1min) and xylene (2×2 min), before being sealed under a coverslip with mounting medium (*Permount*). The cresyl violet solution was prepared beforehand at a concentration of 0.5% in water, filtered and stored at room temperature. For stereological counting of eYFP positive mesencephalic neurons, a second well amongst the six wells containing the mesencephalic sections of each animal was selected at random. The DAB immunostaining protocol against eYFP was performed using a rabbit anti-GFP primary antibody (1:5000) diluted in 0.01M PBS containing 0.3% Triton X-100 and 0.02% NaN_3_, and the same secondary antibody (Biotin-SP goat anti-rabbit at a concentration of 1:200 diluted in 0.01M PBS containing 0.03% Triton X-100, by following the aforementioned steps used for DAB immunostaining against TH, except with a reaction time in DAB reaction solution reduced to 30s due to faster staining.

For stereological counting, a sample of one in six mesencephalic sections containing the SNc/VTA was made, which corresponded to six sections per animal. The SNc and VTA boundaries were manually drawn using the Stereo Investigator software (MBF Bioscience). The *Atlas of the Developing Mouse* Brain (Paxinos et al., 2006) was used as anatomical reference for P15 mice and *The Mouse Brain in Stereotaxic Coordinates* (Franklin & Paxinos, 2008) for P90 mice. The stereological parameters were set to 40μm section thickness, 10μm counting field depth (*optical fractionator*), 50μm by 50μm counting frame, and a 100μm-by-100μm systematic random sampling grid for P15 animals and 150μm by 150μm for P90 animals. A tighter grid was used for P15 animals due to Gundersen M1 coefficients not being inferior to 0.10 (significance threshold value) with a 150μm-by-150μm grid. Also, for all animals Gundersen M1 coefficients were inferior to 0.10 in reference structures (unlesioned structures or structures targeted by the viral transfection), which validated the obtained stereological data. The microscope used for stereological counting was a Leica DMRE using the Stereo Investigator V.9 software and a CX9000 camera (MBF Bioscience). A 10X objective was used for navigation and boundary creation, and a 100X oil immersion objective was used for stereological counting.

### Confocal microscopy image acquisition

In order to evaluate the effect of DA neuron loss on the axonal arborisation of surviving DA neurons, one-in-six of the coronal striatum sections were used. Sections were then immunostained against either TH alone or eYFP and TH, as described previously, and analysed using confocal microscopy at 20X magnification. For each section of the sample, 1 to 6 fields (0.9mm^2^surface per field) were analysed in the right and left dorsal striatum, and 1 to 2 fields were analysed in the right and left ventral striatum, depending on the total surface of the structures on any given section. For a single animal on a given section, fields selected on the right and left structures were equal in numbers and of equivalent anatomical location. For all animals, homologous sections were sampled using the same number of fields situated in homologous anatomical locations. Images were acquired using an Olympus FV1000 confocal microscope and the FV-10 ASW 3.1 software. A 20X water immersion objective was used. For figures, a series of image acquisitions was made using a 60X oil immersion objective. Acquisition parameters were kept constant for section samples that had been immunostained together and comprised equal numbers of subjects from each group, and parameters were determined from the sample having the highest intensity in the group to avoid sensor saturation during image acquisition.

To estimate the volume of DA axonal arborisation in the striatum, images acquired through confocal microscopy for the striatal sampling of an animal were first treated using ImageJ software (NIH, v.1.49r) by converting them to an 8bit format. For each animal group that were simultaneous immunostained and imaged with the same acquisition parameters, a background noise value was determined by calculating an average of maximal signal values for pixels situated outside structures known to express TH (striosomes, corpus callosum, anterior commissure), measured from a random sampling of 10% of images to be analysed. Background noise values obtained for each immunostaining group were subtracted to the signal intensity values of all the images from the corresponding group. From there onwards, a pixel having a signal value higher than 0 was considered to contain fluorescence signal corresponding to the immunostaining target (TH or eYFP), and the surface proportion (*p_n_*) of signal attributed to marked axonal arborizations was calculated for every image *n*. Each structure (ipsilesional dorsal striatum, ipsilesional ventral striatum, contralesional dorsal striatum, contralesional ventral striatum) being evaluated from 1 to 5 images (*n*) per section, the average surface proportion (*x_p_*) of signal was calculated for that structure on a given section. For instance, for a anteroposterior sampling section containing four fields in the left dorsal striatum (*n*=4), four fields in the right dorsal striatum (*n*=4), two fields in the left ventral striatum (*n*=2) and two fields in the right ventral striatum (*n*=2), four *x_p_* averages were calculated using the formula 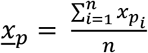. For the four structures, the *x_p_* values were calculated for every section of the sample. To transform these quantifications into a surface (in mm^2^), the area (*a*) of the structures was measured in ImageJ from phase contrast imaging reconstructions of the sections, acquired after completing the acquisitions at the confocal microscope and at the epifluorescence microscope. For each structure, the *x_p_* of a section was multiplied by the total structure area (*a*) on that section to obtain an estimate of the surface (in mm^2^) attributed to signal on that section (*a_s_*). By considering that each coronal section of the sample, cut at a thickness of 40μm, represented the 5 following sections (240μm thickness for 6 sections), the *a_s_* values for each structure were plotted on a chart according to the rostrocaudal position of the corresponding sections on the *x* axis (in mm), and an area under the curve was measured to obtain an estimate of the total volume attributed to signal (*v_s_*) in each structure. By determining the ratio of the TH signal volume (*v_s_*) in the dorsal striatum to the total number of DA neurons in the SNc estimated through stereological counting for each animal, the average volume of signal attributed to a single neuron was calculated for each neuronal group. This value was then used to estimate the average volume of the axonal arborisation of SNc and VTA DA neurons. The same procedure was followed to analyse data obtained with eYFP immunostainings in the P90 DATcre^+/−^ groups transfected with AAV2-eYFP, using images containing the eYFP signal in the striatum and stereological counting results in the mesencephalon for section samples having received DAB immunostaining against eYFP. However, eYFP striatal signal was normalized to the total number of neurons infected regardless of their belonging to the VTA or the SNc.

### Statistical analysis

For every animal group, outlier data detection amongst TH positive stereological neuron counts on the contralesional side were performed using a Grubbs test (extreme studentized deviate), and animals for which outlier data was obtained at a significance threshold of *∝*=0.05 were rejected. This outlier data detection was needed to remove animals with a significantly different number of TH positive neurons in the mesencephalon, either constitutively or secondary to a faulty staining or stereological counting. One animal from the P15 WT group (animal 6) was eliminated from further analysis, because the contralesional SNc contained too few DA neurons. No other animal was rejected from analysis following this criterion. Because the goal of the present study required animals that had undergone a significant DA neuron loss in the SNc, the DA neuron survival ratio in the ipsilesional SNc compared to the contralesional SNc of the same animal was quantified in order to exclude animals that had not undergone any significant lesion in the SNc. A threshold value of 85% DA neuron survival in the ipsilesional SNc following the 6-OHDA injection was considered to exclude animals with too small a lesion. One animal from the P15 DATcre group (animal 14) was rejected under this criterion for the analysis relative to neuronal compensation. For a given animal group, bilateral Student tests paired for ratio were performed for neuronal count values and signal volume values (*vs*) in the striatum, by pairing contralesional and ipsilesional values of each animal. The Student test was parametric (assuming a Gaussian distribution) and paired for ratio. The significance threshold was determined to correspond to a *p*-value *p*<0.05 to reject the null hypothesis of the absence of a difference between contralesional and ipsilesional sides for a given animal.

## Results

### Evidence of compensatory axonal expansion after a neonatal 6-OHDA lesion

Our initial objective was to test the hypothesis that a partial neonatal ablation of SNc DA neurons would lead to adult mice in which a smaller number of SNc DA neurons would each cover a larger striatal territory due to compensatory axonal sprouting. To achieve this, we unilaterally injected postnatal day 5 (P5) DAT-Cre^+/−^ mice in the SNc with a low dose of the DA-neuron-specific toxin 6-OHDA (0.5 μl at 0.25 mg/ml). After 10 days, mice were sacrificed, DA neuron populations were estimated using unbiased stereological counting and the density of TH positive axonal varicosities in the dorsal and ventral sectors of the striatum was quantified using confocal microscopy. We found that in DAT-Cre^+/−^ mice, this low dose of 6-OHDA caused a significant loss of SNc DA neurons on the lesioned side (LS) relative to the control side (CS) (40.26%, p = 0.0163; LS = 3722 ± 570.1 neurons, CS = 6230 ± 518.6 neurons) (**Fig. 1A and E**) and a significant but proportionally smaller decrease in the volume of TH-positive processes in the dorsal striatum on the lesioned side (22.64%, p = 0.047; LS = 13.43 ± 0.9232 mm^3^, CS = 17.36 ± 1.36 mm^3^) (**Fig. 1B, D and F**), thus reflecting a rapid and efficient compensation of the axonal domain. This first experiment did not lead to a significant loss of DA neurons in the closely located ventral tegmental area (VTA) (p = 0.4768, LS = 8894 ± 855.8 neurons, CS = 9497 ± 807.4 neurons) (**Fig. 1A and E**) or to a significant decrease in the volume of DA axonal varicosities in the ventral striatum (p = 0.236, LS = 2.878 ± 0.2831 mm^3^, CS = 3.014 ± 0.2841 mm^3^) (**Fig. 1C and F**). Comparing the relative change in axonal arbor size, obtained by normalizing striatal TH signal to the number of DA neurons, suggests that TH positive axonal arbors in the dorsal striatum, normally associated with SNc DA neurons, significantly increased in size (38.42% increase in size, *p* = 0.0111; LS = 4.003 ± 0.5616 × 10^6^ μm^3^, CS = 2.892 ± 0.3759 × 10^6^ μm^3^) (**Fig. 1G**). The TH positive axonal arbor in the ventral striatum, associated with VTA DA neurons, did not undergo significant expansion (p = 0.7695, LS = 0.3276 ± 0.02056 × 10^6^μm^3^, CS = 0.3225 ± 0.02503 × 10^6^μm^3^) (**Fig. 1G**).

**Figure 1:**
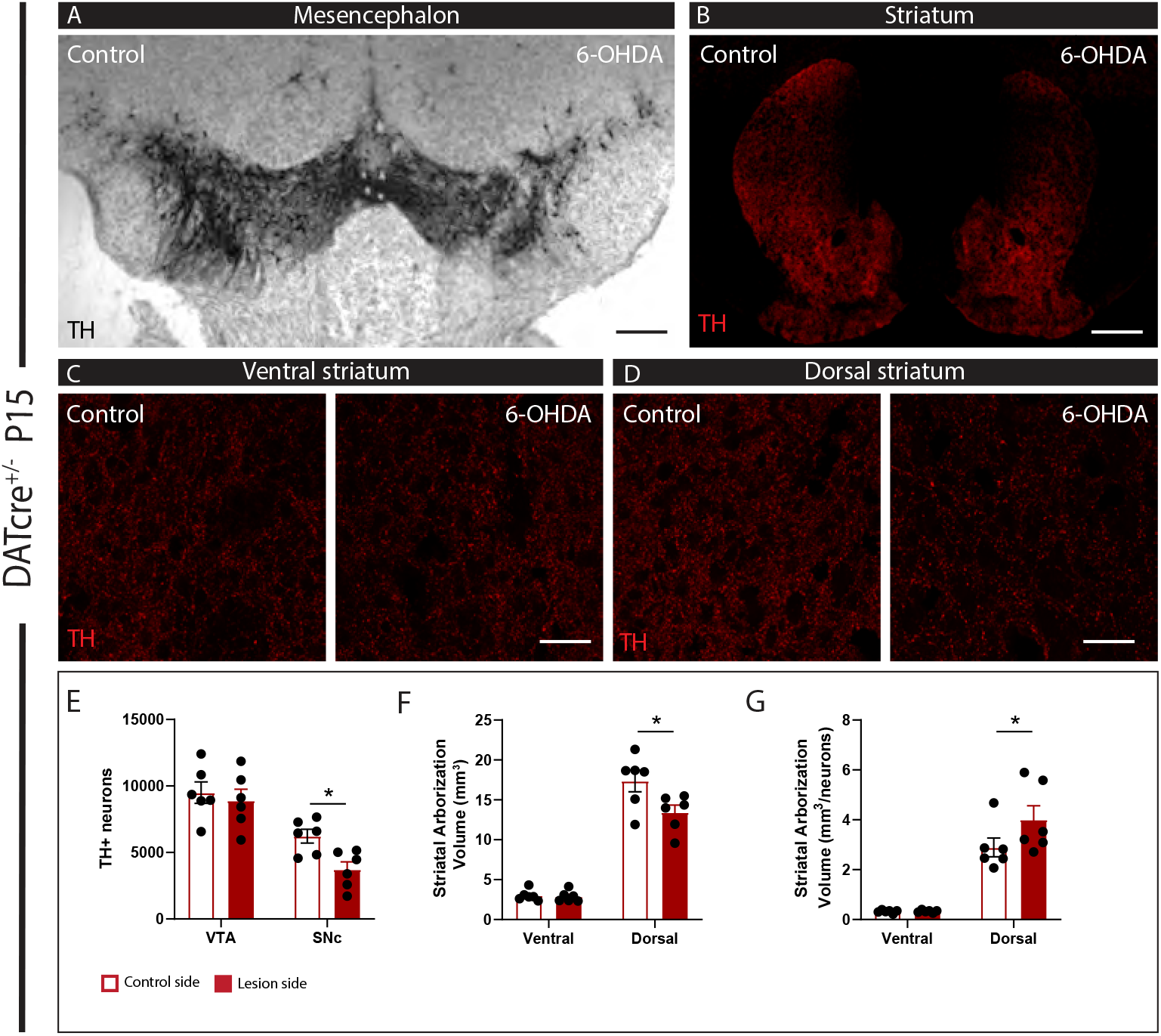
Increased TH positive axonal arborization volume in the dorsal striatum from surviving DA neurons 10 days following partial neonatal 6-OHDA lesion of the SNc. (**A-B**) TH immunohistochemistry showing a reduced density of TH positive neurons in the SNc and a small reduction of the density of TH positive axonal fibers in the dorsal striatum on the lesioned (right) side of the brain. Scale bar = 1 mm for panel A and 500 μm for panel B. (**C-D**) Higher power magnification showing TH positive axonal varicosities in the ventral and dorsal striatum. Scale bar= 40 μm. (**E**) Stereological counting of the number of DA neurons in the SNc and VTA on the control and 6-OHDA sides. (**F**) Quantification of the relative density of TH positive axonal varicosities in the ventral and dorsal striatum obtained from confocal images. (**G**) Estimated axonal arbor volume DA neurons projecting to the ventral or dorsal striatum obtained by normalizing TH positive striatal volumes by the number of associated SNc or VTA DA neurons. Data are presented as mean +/− standard error of the mean (SEM), N = 6 animals/group.

A separate cohort of mice underwent the same unilateral intra-nigral 6-OHDA injection at P5, but were sacrificed and examined 85 days later, at P90. Here again, the 6-OHDA lesion caused a significant decrease in the number of SNc DA neurons (44.34%, p < 0.0001; LS = 3585 ± 339.2 neurons, CS = 6441 ± 362.5 neurons) (**Fig. 2A and E**). However, the associated volume loss of TH positive processes in the dorsal striatum observed at P15 was no longer visible in these mice at P90 (p = 0.1224, LS = 11.24 ± 0.9199 mm^3^, CS = 12.24 ± 0.5762 mm^3^) (**Fig. 2B, D and F**). Quantification of axonal varicosity density in the dorsal striatum, normalized to the number of associated DA neurons in the SNc, confirmed that robust compensatory axonal sprouting was maintained until P90 (68.18% increase in size, p < 0.0001; LS = 3.256 ± 0.2787 × 10^6^μm^3^, CS = 1.936 0.1166 × 10^6^μm^3^) (**Fig. 2G**). In this group, there was a small but significant reduction in the number of VTA DA neurons (14.36%, p = 0.0044; LS = 6974 ± 387.7 neurons, CS = 8143 ± 446 neurons) and no significant change in the measured volume of DA varicosities in the ventral striatum (p = 0.3902, LS = 1.679 ± 0.1188 mm^3^, CS = 1.733 ± 0.09834 mm^3^). Quantification of axonal varicosity density in the ventral striatum, normalized to the number of associated DA neurons in the VTA did not reveal a significant change, suggesting that VTA DA neurons did not undergo any major compensatory sprouting under such conditions and in the presence of a moderate SNc lesion (p = 0.1399, LS = 0.248 ± 0.02534 × 10^6^μm^3^, CS = 0.219 ± 0.0195 × 10^6^μm^3^) (**Fig. 2E, F and G**).

**Figure 2:**
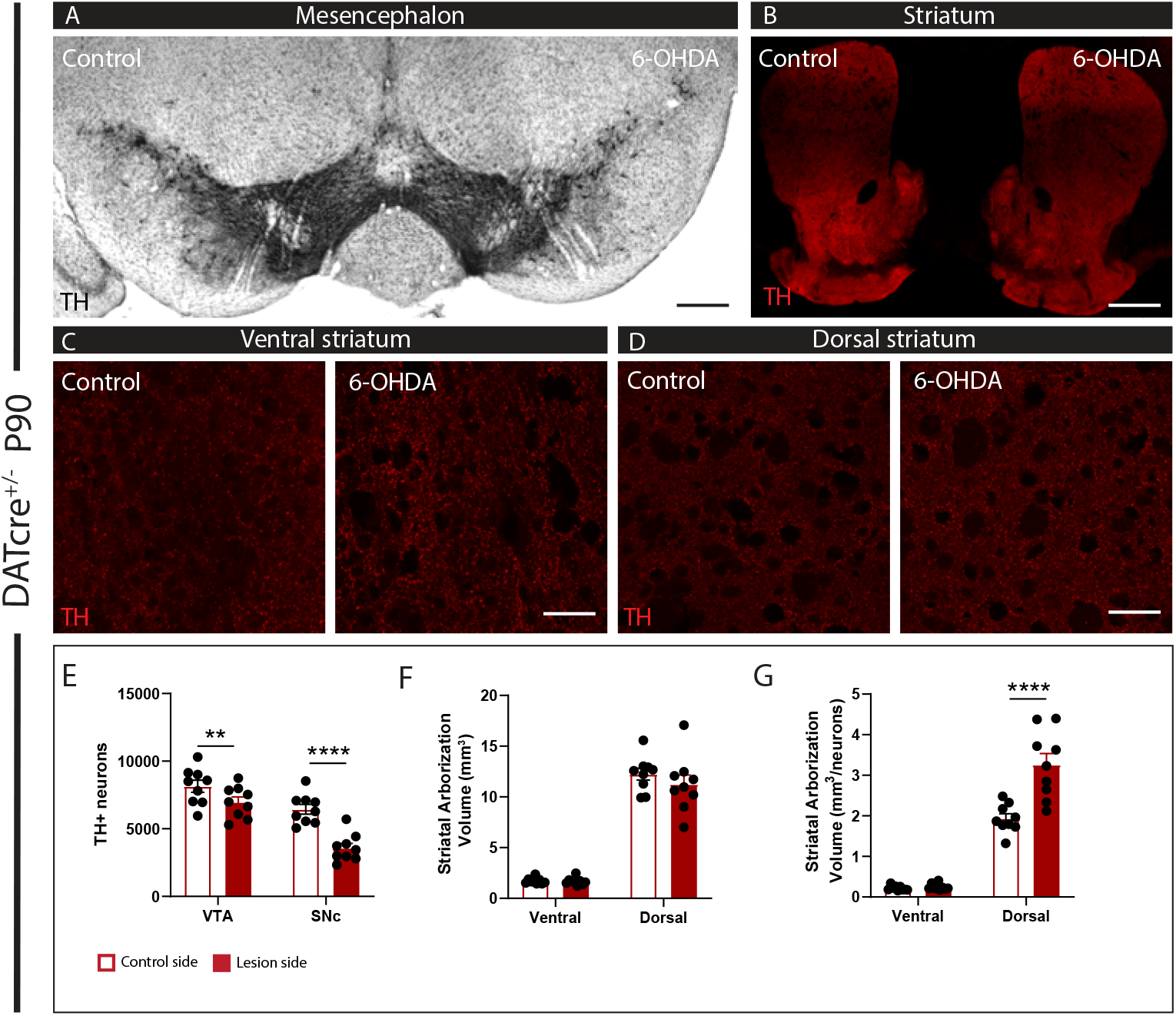
Increased TH positive axonal arborization volume in the dorsal striatum from surviving DA neurons eighty-five days following partial neonatal 6-OHDA lesion of the SNc. (**A-B**) TH immunohistochemistry showing a reduced density of TH positive neurons in the SNc and the absence of notable reduction of the density of TH positive axonal fibers in the dorsal striatum on the lesioned (right) side of the brain. Scale bar = 1 mm for panel A and 500 μm for panel B. (**C-D**) Higher power magnification showing TH positive axonal varicosities in the ventral and dorsal striatum. Scale = 40 μm. (**E**) Stereological counting of the number of DA neurons in the SNc and VTA on the control and 6-OHDA sides. (**F**) Quantification of the relative density of TH positive axonal varicosities in the ventral and dorsal striatum obtained from confocal images. (**G**) Estimated axonal arbor volume DA neurons projecting to the ventral or dorsal striatum obtained by normalizing TH positive striatal volumes by the number of associated SNc or VTA DA neurons. Data are presented as mean +/− standard error of the mean (SEM), N = 9 animals/group.

### Viral mediated labelling confirms compensatory axonal expansion in SNc DA neurons after a neonatal 6-OHDA lesion and suggests little if any contribution of VTA DA neurons

Although it is well established that VTA DA neurons projects mainly to the ventral striatum with only limited axons reaching the various sectors of the dorsal striatum, we next aimed to directly visualize the striatal projections of SNc and VTA neurons separately in this same neonatal lesion model. We hypothesized that the axonal projections of SNc DA neurons in the dorsal striatum should expand after the lesion. Furthermore, we hypothesized that VTA DA neurons should show more limited changes in their axonal arbor density. For this, we took advantage of a floxed YFP AAV construct injected stereotaxically in the SNc or VTA of DAT-Cre^+/−^ mice 55 days after the P5 6-OHDA injection (thus at P60). In a first group of mice, a small volume of 0,8 μl of virus was injected in the SNc on both the lesioned and non lesioned sides of the brain (**Fig. 3A**). In a separate group of mice, a smaller volume of 0,4 μl of virus was injected centrally in the VTA to target VTA neurons on both sides of the brain (**Fig. 4A**). Mice were sacrificed 30 days later at P90 to quantify the number of infected DA neurons in either the SNc or the VTA, as well as the density of YFP-positive axonal varicosities in the dorsal and ventral striatum.

**Figure 3:**
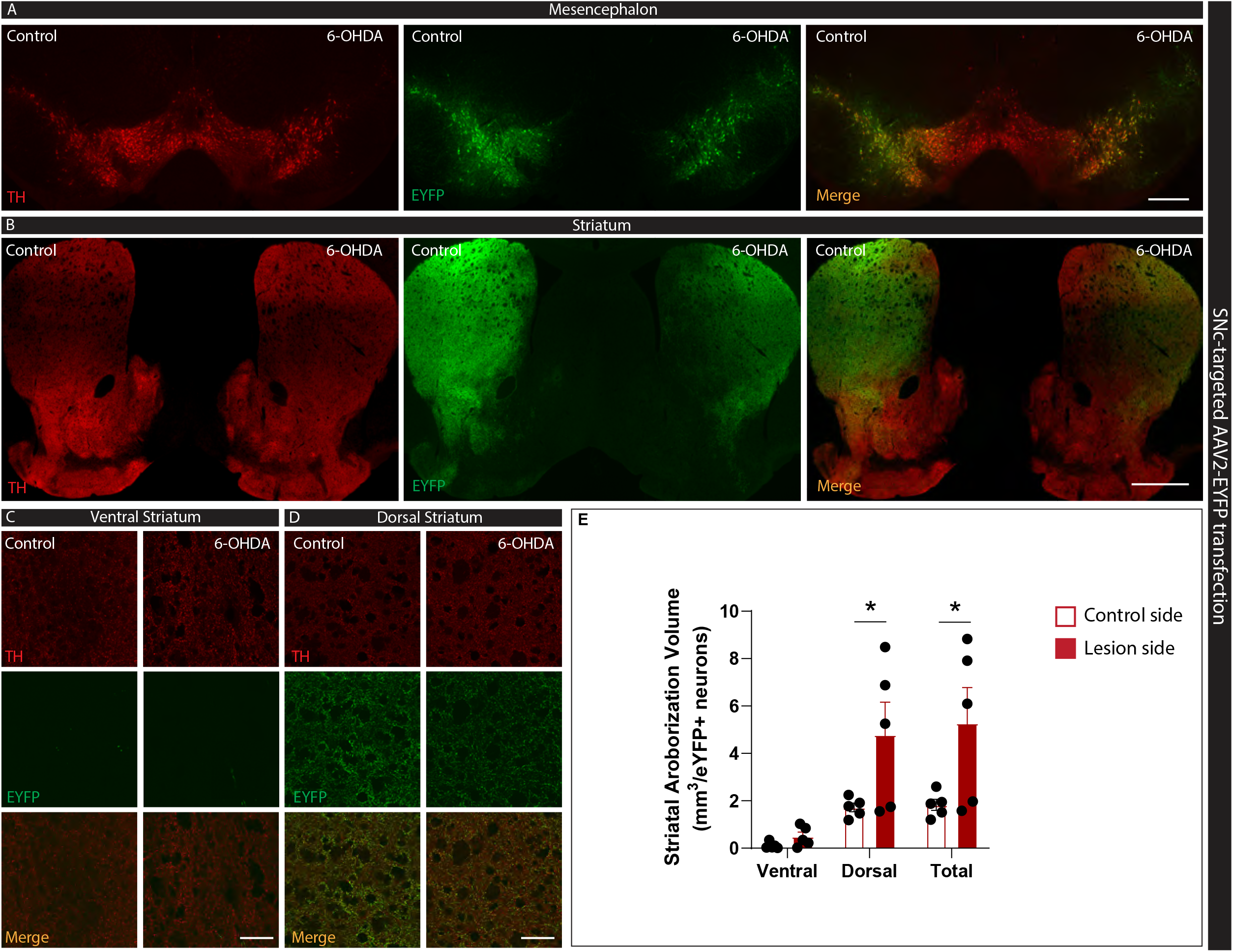
Surviving SNc DA neurons show robust compensatory axonal sprouting in the dorsal striatum following partial 6-OHDA lesion of the SNc at P5. Injection of a floxed eYFP expressing AAV in the SNc of DAT-Cre mice was used to specifically label SNc neurons and quantify their axonal arborization size in the striatum following partial neonatal 6-OHDA lesion of the SNc. (**A-B)** TH (red) and eYFP (green) immunohistochemistry illustrating labelling of SNc DA neuron axonal projections, predominantly in the dorsal striatum. The merge images show that most infected neurons were localized in the SNc, but a small subset of lateral VTA DA neurons were also infected. Scale bar = 1 mm for panel A and 500 μm for panel B. (**C-D**) Higher power magnification showing TH and EGFP-labelled DA axonal varicosities in the ventral and dorsal sectors of the striatum. Very few eYFP fibers were detected in the ventral striatum, confirming that most infected neurons were from the SNc or from lateral VTA neurons that mostly project to the dorsal striatum. Scale = 40 μm. (**E**) Estimate of relative axonal arbor volume of SNc DA neurons. Data are presented as mean +/− standard error of the mean (SEM), N = 5 animals/group.

**Figure 4:**
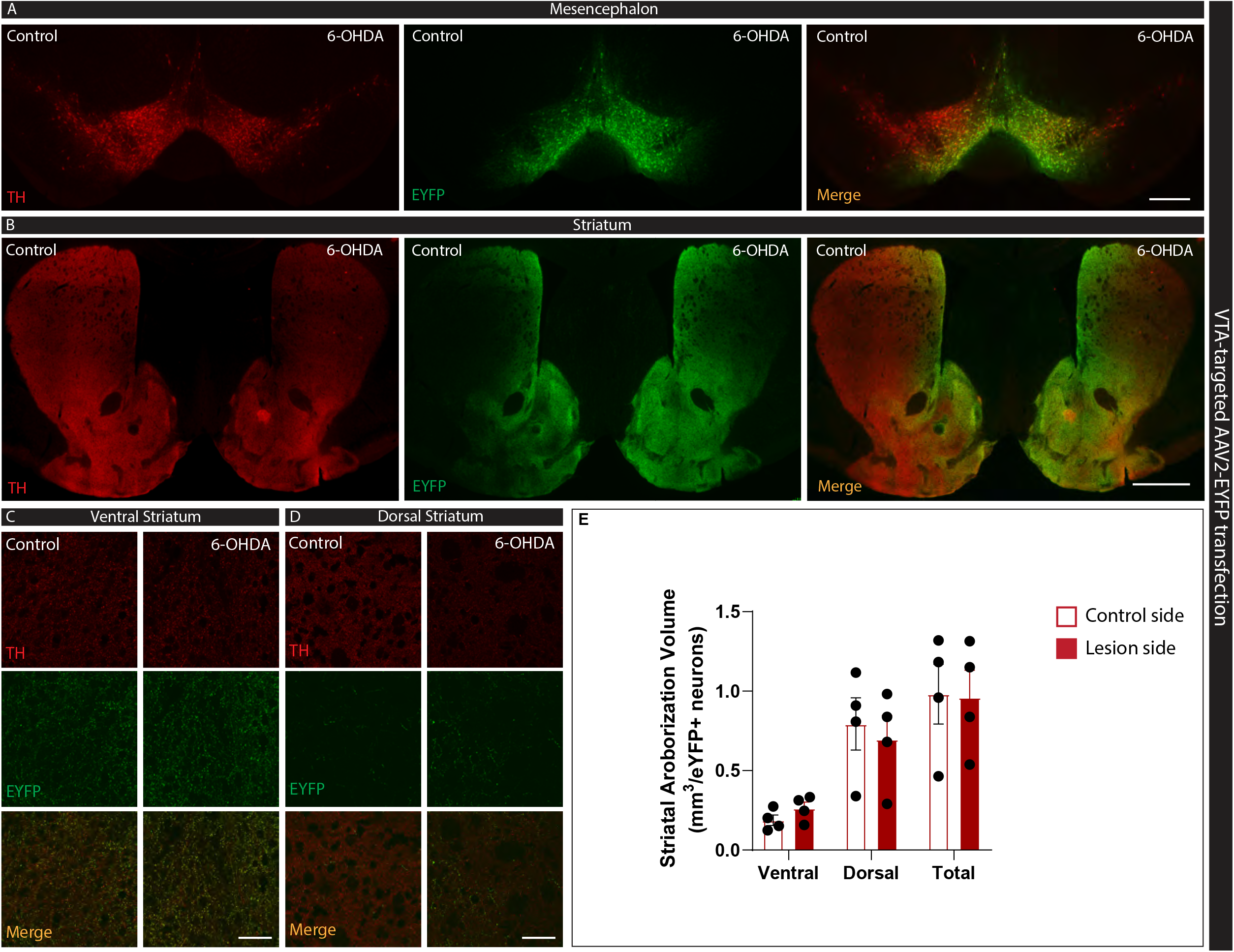
No compensatory axonal sprouting from VTA DA neurons following partial 6-OHDA lesion of the SNc at P5. Injection of a floxed eYFP expressing AAV in the VTA of DAT-Cre mice was used to specifically label VTA neurons and quantify their axonal arborization size in the striatum following partial neonatal 6-OHDA lesion of the SNc. (**A-B)** TH (red) and eYFP (green) immunohistochemistry illustrating labelling of VTA DA neuron axonal projections, predominantly in the ventral striatum and median dorsal striatum, closer to the midline. The merge images show that most infected neurons were localized in the VTA, but a small subset of medial SNc DA neurons were also infected. Scale bar = 1 mm for panel A and 500 μm for panel B. (**C-** Higher power magnification showing TH and eYFP-labelled DA axonal varicosities in the ventral and dorsal sectors of the striatum. Fewer eYFP fibers were detected in the dorsal striatum, confirming that most infected neurons were from the VTA neurons that mostly project to the ventral striatum. Scale = 40 μm (**E**) Estimate of relative axonal arbor volume of VTA DA neurons. Data are presented as mean +/− standard error of the mean (SEM), N = 4 animals/group.

For experiments aiming to label SNc DA neurons, although the number of transfected DA neurons varied substantially between different mice, a majority of the transfected DA neurons were found in the SNc (CS = 63.95 % and LS = 84.84%). Since a variable subset of VTA neurons were also infected when targeting the SNc, in some mice, a modest subset of the YFP labelled terminals in the striatum could potentially belong to VTA DA neurons. However, the density of YFP positive axonal varicosities detected in the ventral striatum was systematically quite low, while it was dense in the dorsal striatum (**Fig. 3B, C and D**). Quantification of the relative density of YFP positive axonal varicosities in the striatum, normalized to the total number of infected DA neurons, revealed that on the lesioned side, the axonal arbor volume of infected neurons was larger compared to the non lesioned side in the dorsal striatum (2.8-fold increase, p = 0.0497, CS = 1.720 ±0.1822 × 10^6^ μm^3^, LS = 4.785 ± 1.378 × 10^6^ μm^3^), but not in the ventral striatum (p = 0.9843, CS = 0.106 ± 0.0625 × 10^6^ μm^3^, LS = 0.492 ± 0.1912 × 10^6^ μm^3^) (**Fig. 3E**). These results thus provide further support for our hypothesis proposing that neonatal ablation of SNc DA neurons leads to early and sustained expansion of their axonal arbor volume in the dorsal striatum of adult mice.

In experiments aiming to label the VTA DA neurons, we found that the vast majority of transfected neurons were within the VTA, with few SNc DA neurons infected (CS = 85.85% and LS = 94.31%) (**Fig. 4A**). Both on the lesioned and non-lesioned sides of the brain, infected VTA neurons projected preferentially to the ventral sectors of the striatum as well as to the medial part of the dorsal striatum, close to the midline (**Fig. 4B**). Quantitative analysis revealed that the normalized global density of YFP labelled axonal varicosities in both the dorsal and ventral striatum was unchanged on the lesioned compared to the non lesioned side of the brain (p = 0.9754, CS = 0.188 ± 0.03291 × 10^6^ μm^3^, LS = 0.263 ± 0.03945 × 10^6^ μm^3^ for ventral striatum and p = 0.9501, 0.794 ± 0.1644 × 10^6^ μm^3^, LS = 0.697 ± 0.1490 × 10^6^ μm^3^ for dorsal striatum) (**Fig. 4C, D, E**). The absence of any significant compensatory axonal sprouting by VTA DA neurons is in line with the fact that the 6-OHDA lesion in these mice mostly spared VTA DA neurons and their projections to the ventral striatum.

## Discussion

The present results provide strong support for the hypothesis that early neonatal ablation of a subset of SNc DA neurons represents a good strategy to force the remaining SNc DA neurons to expand their axonal arbor in the dorsal striatum. We found that following ablation of approximately half of DA neurons at P5, the surviving SNc DA neurons show close to a 3-fold increase in the global volume of their axonal arbor size in the dorsal striatum, a process that is already well advanced as soon as 10 days following the lesion. Furthermore, examination of the respective projections of SNc and VTA DA neurons revealed that SNc DA neurons are likely to account for the bulk of this compensatory axonal sprouting.

Our finding of extensive axonal sprouting by SNc DA neurons is in line with previous results demonstrating that mesencephalic DA neurons can activate homeostatic mechanisms leading to partial recovery of DAergic axonal projections in the dorsal striatum after partial lesions in adult rodents (Blanchard et al., 1996). In the adult rat brain, it has been suggested that lesion sizes lower than 80% were associated with only modest compensatory axonal growth in the striatum, whereas compensatory sprouting, associated with upregulation of GAP-43 mRNA, an axonal growth marker, occurs with DA depletion levels of 95% (Robinson et al., 1994; Song et al., 2000). The molecular mechanisms triggering this compensatory axonal sprouting are ill-defined, but local inflammatory signals mediated by cytokines secreted by astrocytes and microglia have been proposed to play a role. Ho and coll. demonstrated that a MPTP lesion induces synthesis of interleukin-1 (IL-1), a cytokine promoting DA neuron axonal sprouting when administered exogenously in the lesioned striatum of mice, which was most significant in young adult mice (2 months) and happened both in the ventral and dorsal striatum, whereas in older mice (8 months), the increase in IL-1 levels was much weaker and limited to the dorsal striatum (Ho and Blum, 1998). This suggests an increased capacity for compensatory DA neuron axonal sprouting throughout the striatum in younger animals. Growth factors such as GDNF and BDNF have also been suggested to contribute to such sprouting (Ho and Blum, 1997; Rosenblad et al., 1998, 2000).

Post-lesional compensatory axonal sprouting has also been previously investigated in neonatal rodents after administration of 6-OHDA (Fernandes Xavier et al., 1994, Luthman et al., 1990, Luthman et al., 1987, Sivam et al., 1987, Snyder et al., 1986, Stachowiak et al., 1984, Stellar et al., 1988, and Towle et al., 1989). However, doses of 6-OHDA leading to near complete loss of SNc DA neurons were used in these previous studies, and as such, the compensatory mechanisms activated in SNc neurons were not studied. The present work is thus the first to examine axonal compensatory sprouting of SNc DA neurons after an early post-natal lesion.

Our comparison of changes in TH immunoreactivity in the ventral and dorsal striatum after partial SNc lesions showed no significant change in the ventral striatum, suggesting that SNc DA neurons, known to preferentially project to the dorsal striatum, contributed preferentially to the compensatory increase detected in the dorsal striatum. However, because VTA neurons can also send some of their branches to sectors of the dorsal striatum, the contribution of VTA neurons to the observed restoration of TH signal in the dorsal striatum was impossible to evaluate in our first series of experiments. Our second series of experiments, using conditional expression of YFP in DA neurons, confirmed that VTA neurons did not contribute significantly to compensatory reinnervation of the dorsal striatum in the present neonatal lesion model.

Our comparison of relative axonal arbor density in the present study was primarily aimed at evaluating the impact of partial lesions on axonal sprouting. However, we were able to also compare axonal arbor volume between SNc and VTA DA neurons. Comparing the data shown in panels G of figures 3 and 4, it is apparent that the relative axonal arbor size of SNc DA neurons is multiple fold larger compared to that of VTA DA neurons. This result is in line with a previous *in vitro* comparison of VTA and SNc DA neurons (Pacelli et al., 2015) and with recent *in vivo* work comparing axonal arbor size of these two populations of DA neurons and the impact of D2 autoreceptors in regulating this morphological characteristic of DA neurons (Giguère et al., 2019). Excitingly, this later report showed that expansion of the axonal arbor of SNc DA neurons caused by conditional genetic deletion of the D2 autoreceptor in DA neurons increased DA neuron vulnerability to 6-OHDA. Such results make it reasonable to propose that in mice with reduced numbers of SNc DA neurons such as described in the present work, subsequent vulnerability to various neurotoxic insults at the adult stage could be increased compared to control animals. Additional work will be requited to test this hypothesis.

In the present work, we used a low dose of 6-OHDA (0.5 μl of a 0.25 mg/ml solution) to induce degeneration of SNc DA neurons. This represents a very modest dose compared to what is usually administered to adult mice or rats. The present experiments were performed using DATcre^+/−^ mice, a knockin line in which one of the copies of the DAT gene locus was replaced with the Cre recombinase sequence. Because these mice therefore have only 50% of the normal levels of DAT protein and 6-OHDA enters DA neurons through DAT, it is possible that in experiments performed in other lines of mice with normal DAT levels, more extensive loss of SNc DA neurons would be obtained with the dose used here in DATcre^+/−^ mice (0.5 μl at 0.25 μg/μl). However, in preliminary experiments performed with wild-type mice with the same dose did not reveal a greatly elevated level of SNc cell loss but importantly replicated our observation of compensatory increase in the density of TH positive axonal processes in the dorsal striatum following this type of partial lesion (**Supplementary figures 1 and 2**).

In conclusion, the present work paves the way to further experiments to better determine the relationship between axonal arbor size and neuronal vulnerability in PD by validating a model allowing the expansion of mesencephalic DA neuron axonal arbor size. It also confirms that neonatal SNc DA neurons have the capacity to increase their axonal arborization size to compensate for the loss of neighboring DA neurons. Additional experiments exploring a broader spectrum of lesion severity will also be needed to improve characterisation of the relationship between lesion size and compensatory axonal growth in neonate mice.

## Acknowledgments

This work was funded by a grant from the Canadian Institutes of Health Research (CIHR), by the Krembil and Brain Canada Foundations, as well as by a pilot project grant from Parkinson Society Canada. Support to the Trudeau lab from the Henry and Berenice Kaufmann Foundation is also acknowledged. Charles Ducrot was supported by a Parkinson Canada studentship.

