## Supplementary figures for "Neonatal 6-OHDA lesion of the SNc induces striatal compensatory sprouting from surviving SNc dopaminergic neurons without VTA contribution"

Fig.S1

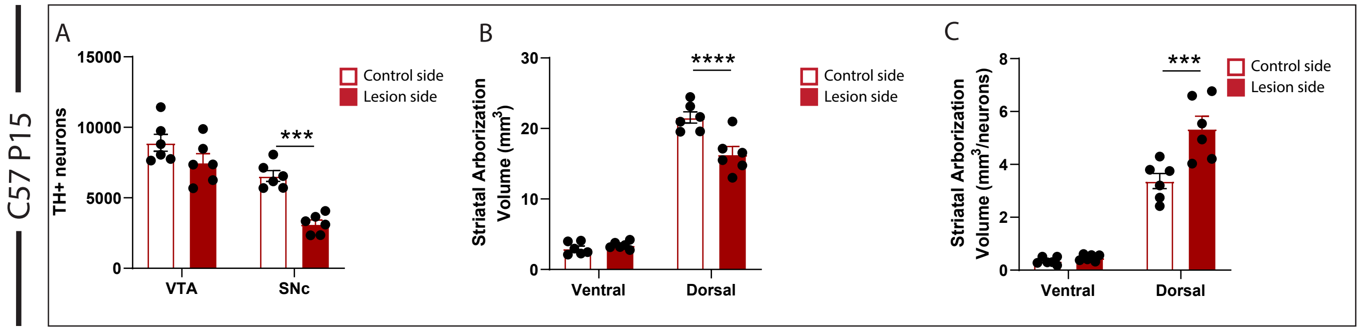
C57 P15
A
TH+ neurons
15000
10000
5000
0
VTA
SNc
***
Control side
Lesion side
B
Striatal Arborization Volume (mm3)
30
20
10
0
Ventral
Dorsal
****
Control side
Lesion side
C
Striatal Arborization Volume (mm3/neurons)
8
6
4
2
0
Ventral
Dorsal
***
Control side
Lesion side

**Supplementary Figure 1 legend: Increased TH positive axonal arborization volume in the dorsal striatum from surviving DA neurons 10 days following partial neonatal 6-OHDA lesion of the SNc of C57/BL6 mice.** (**A**) Stereological counting of the number of DA neurons in the SNc and VTA on the control and 6-OHDA sides. (**B**) Quantification of the relative density of TH positive axonal varicosities in the ventral and dorsal striatum obtained from confocal images. (**C**) Estimated axonal arbor volume DA neurons projecting to the ventral or dorsal striatum obtained by normalizing TH positive striatal volumes by the number of associated SNc or VTA DA neurons. Data are presented as mean +/- standard error of the mean (SEM), N = 6 animals/group.

### Fig.S2

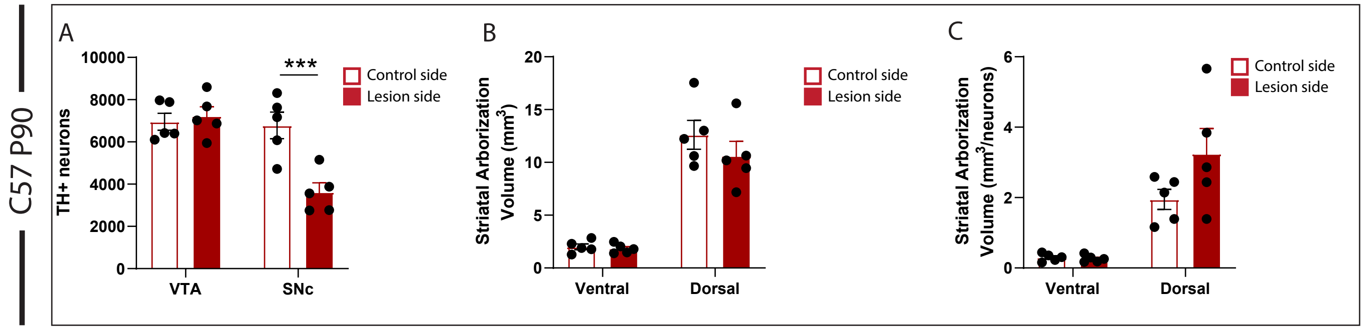
C57 P90
A
TH+ neurons
10000
8000
6000
4000
2000
0
VTA
SNc
***
Control side
Lesion side
B
Striatal Arborization Volume (mm3)
20
15
10
5
0
Ventral
Dorsal
Control side
Lesion side
C
Striatal Arborization Volume (mm3/neurons)
6
4
2
0
Ventral
Dorsal
Control side
Lesion side

**Supplementary Figure 2 legend: Increased TH positive axonal arborization volume in the dorsal striatum from surviving DA neurons eighty-five days following partial neonatal 6-OHDA lesion of the SNc.** (**A**) Stereological counting of the number of DA neurons in the SNc and VTA on the control and 6-OHDA sides. (**B**) Quantification of the relative density of TH positive axonal varicosities in the ventral and dorsal striatum obtained from confocal images. (**C**) Estimated axonal arbor volume DA neurons projecting to the ventral or dorsal striatum obtained by normalizing TH positive striatal volumes by the number of associated SNc or VTA DA neurons. Data are presented as mean +/- standard error of the mean (SEM), N = 5 animals/group.
